# A maternal-to-zygotic-transition gene block on the zebrafish sex chromosome

**DOI:** 10.1101/2023.12.06.570431

**Authors:** Catherine A. Wilson, John H. Postlethwait

## Abstract

Wild zebrafish (*Danio rerio*) have a ZZ/ZW chromosomal sex determination system with the major sex locus on the right arm of chromosome-4 (Chr4R) near the largest heterochromatic block in the genome, suggesting the hypothesis that the Chr4R transcriptome might be different from the rest of the genome. We conducted an RNA-seq analysis of adult ZW ovaries and ZZ testes and identified four regions of Chr4 with different gene expression profiles. Unique in the genome, protein-coding genes in a 41.7 Mb section (Region-2) were expressed in testis but silent in ovary. The AB lab strain, which lacks sex chromosomes, verified this result, showing that testis-biased gene expression in Region-2 depends on gonad biology, not on sex-determining mechanism. RNA-seq analyses in female and male brain and liver validated few transcripts from Region-2 in somatic cells, but without sex-specificity. Region-2 corresponds to the heterochromatic portion of Chr4R and its content of genes and repetitive elements distinguishes it from the rest of the genome. In Region-2, protein-coding genes lack human orthologs; it has zinc finger genes expressed early in zygotic genome activation; it has maternal 5S rRNA genes, maternal spliceosome genes, a concentration of tRNA genes, and an distinct set of repetitive elements. The colocalization of 1) genes silenced in ovaries but not in testes that are 2) expressed in embryos briefly at the onset of zygotic genome activation; 3) maternal-specific genes for translation machinery; 4) maternal-specific spliceosome components; and 4) adjacent genes encoding miR-430, which mediates maternal transcript degradation, suggest that this is a Maternal-to-Zygotic-Transition Gene Regulatory Block.

**ARTICLE SUMMARY:** The wild zebrafish sex chromosome has a region, unique in the genome, that contains protein-coding genes silenced in ovaries but expressed in testes and transiently in the embryo as it begins to express its own genes. This region also contains maternal-specific genes encoding the protein-synthesis machinery used specifically by developing embryos, and molecules that target for degradation messenger RNAs that the mother stored in her eggs. This region defines a distinct maternal-to-zygotic-transition gene block.

## INTRODUCTION

Within a species, selection can act differently on females and males due to differences in physiological traits that affect gamete formation, mating, fertilization, and parental care (Darwin 1871). In humans, these sex-related physiological features can lead to health-related sex-biased discrepancies in functioning of the immune, cardiovascular, and skeletal systems, and in metabolism, responses to drugs and toxins, and the types and incidences of cancers (Wizemann and Pardue 2001). These facts led some health research funding agencies to prioritize research that balances considerations of sex as a biological variable (Clayton and Collins 2014). Sex differences in development and physiology can arise from genes that are expressed at a greater level in one sex compared to the other (sex-biased genes) (Dutoit et al. 2018).

Genes with sex-biased expression might not be distributed randomly across the genome. In species that have chromosomally XX females and XY males, X chromosomes are often enriched with female-biased genes and conversely, in species that are ZW females and ZZ males, Z chromosomes often have an excess of male-biased genes (Mank 2009). These distributions likely reflect the fact that the phenotypes of traits that maximize male fitness might sometimes conflict with those that maximize female fitness (Lande 1980).

Fish make superior models for understanding sex genetics, gonad development, and the mechanisms of human diseases (Henke et al. 2022; MacRae and Peterson 2022; Aharon and Marlow 2021; Ye and Chen 2020; Nagahama et al. 2021; Pan et al. 2021; Baldridge et al. 2021) and sex-biased expression is common in fish for protein-coding, microRNA (miRNA), and long non-coding RNA (lncRNA) genes (Desvignes et al. 2019) (Desvignes et al. 2021; Dechaud et al. 2021; Guo et al. 2021; Lichilin et al. 2021; Kaitetzidou et al. 2022). In zebrafish, a premier model for human disease (Baldridge et al. 2021), sex-biased transcription has been studied for liver, brain, and gonads, with by far the greatest sex differences in gene expression found in the gonads (Robison et al. 2008; Sreenivasan et al. 2008; Small et al. 2009; Ung et al. 2013; Zheng et al. 2013; Wong et al. 2014; Yang et al. 2016). Furthermore, in zebrafish and other teleosts, female-biased and male-biased genes both evolve more rapidly than genes that don’t show sex-biased expression (Yang et al. 2016; Lichilin et al. 2021).

Despite the importance of sex-biased gene expression to help understand sexual selection, to discover the origin of sex-related phenotypes, and to evaluate the accumulation of sex-related genes on sex chromosomes, little information exists concerning the genomic distribution of sex-biased genes along fish chromosomes (Wong et al. 2014; Kaitetzidou et al. 2022) . Here, we describe the discovery of strong testis-biased gene expression in hundreds of protein-coding genes in a heterochromatic region of the long arm of chromosome-4 near the sex-determining locus in a zebrafish strain that retains the wild ZW female / ZZ male sex determining system (Wilson et al. 2014). Results showed that this region coincides with genes encoding maternally supplied ribosomal and spliceosomal components for the early embryo (Locati et al. 2017a; Pagano et al. 2020b) as well as genes that are among the first to be transcribed after zygotic gene activation (White et al. 2017), and furthermore, is adjacent in the genome to genes that help in the removal of maternal transcripts (Giraldez et al. 2005; Takacs and Giraldez 2016; Giraldez et al. 2006; Lund et al. 2009), making it a Maternal-to-Zygotic Transition Gene Block.

## METHODS

Female and male zebrafish of the Nadia strain (NA, ZDB-GENO-030115-2) provided gonads for these experiments. All animal work was approved by the University of Oregon Institutional Animal Care and Use Committee (# AUP-21-06 v.1). Adult fish were euthanized at 3-months post-fertilization and genotyped by PCR for sex chromosomes as previously described (Wilson et al. 2014). From five adult ZW females and five adult ZZ males, we dissected pairs of ovaries and testes, respectively, stored them in RNAlater (Invitrogen), and extracted total RNA using the RiboPure RNA Purification Kit (Ambion). We assessed total RNA quality on the Agilent Fragment Analyzer and enriched mRNA from samples with an RQN ≥ 7.9 using the Dynabeads mRNA Purification Kit (Ambion). Strand-specific sequencing libraries were generated with the NextFlex Rapid Directional qRNA-Seq Library Prep Kit (BIOO Scientific) and were sequenced on an Illumina NextSeq 500 to generate paired-end 75-nt reads. Illumina reads were pre-processed with Dupligänger (Sydes et al. 2019) to annotate barcodes, to act as a wrapper for Cutadapt (Martin 2011), which removes adapters, and Trimmomatic (Bolger et al. 2014), which trims reads for quality, and to remove PCR duplicates. Reads were then aligned to the zebrafish reference genome (version GRCz11) (Howe et al. 2013) with GSNAP (Wu and Nacu 2010), allowing up to 10% mismatches and using transcript splice sites identified in the Ensembl Release 92 annotation (Ens92) (Zerbino et al. 2018). Reads that mapped to annotated features were counted with HTSeq-Count in ‘strict’ mode (Anders et al. 2015). Differential expression in protein-coding genes was calculated with DESeq2 (version1.20.0) (Love et al. 2014). We performed functional enrichment analysis of differentially expressed genes using PANTHER (Mi et al. 2017).

RNA-seq data from zebrafish strain AB (ZDB-GENO-960809-7, derived by multiple rounds of gynogenesis) wild-type ovary and testis were obtained from SRA accession PRJNA512103 and processed as described (Yan et al. 2019). AB control liver and brain RNA-seq data were obtained from SRA accession PRJNA600413 (Banu et al. 2020). These libraries did not have molecular barcodes, so we removed the adapters with Cutadapt (Martin 2011), quality trimmed with Trimmomatic (Bolger et al. 2014), and then aligned to GRCz11 as above. We used GTFtools (Li et al. 2022) to analyze Ens92 gene models and used the length of merged exons of isoforms of each gene to calculate TPM (transcripts per kilobase of gene length per million total reads) for all protein-coding genes in each library.

The GRCz11 repeat annotation was obtained from UCSC (Navarro Gonzalez et al. 2021). We excluded any repeat with a “Family” or “Class” that contained a “?”, or was labeled “Unknown”. We also excluded any repeat with the following “Family” or “Class” : “Simple_repeat”, “Low_complexity”, “tRNA”, “rRNA”, and “snRNA” as well as repeats with fewer than 50 copies in the genome. We generated bed files for each repeat name and used BEDTools (Quinlan and Hall 2010) to calculate the overlap with selected genomic regions, then calculated the percentage of each region covered by each repeat type. Hierarchical clustering of genomic regions was performed using the R function “hclust” and the “average” method. Principal component analysis based on the correlation matrix was performed using the R function “prcomp”. Transposable element transcription quantification was calculated using Telescope (Bendall et al. 2019). Reads from Nadia gonads were realigned to the GRCz11 zebrafish assembly using STAR (Dobin et al. 2013) with parameters --winAnchorMultimapNmax 100 -- outFilterMultimapNmax 100. Alignments were then reassigned with Telescope, using the UCSC GRCz11 repeat annotation as input. TE counts generated by Telescope were analyzed using DESeq2 (Love et al. 2014).

The zebrafish maternal 5S rDNA consensus sequence 5Sseq1 (Locati et al. 2017b) was aligned to GRCz11 with BLASTN (Camacho et al. 2009) using parameters -perc_identity 100 - qcov_hsp_perc 100. Maternal 5S gene positions were plotted on Chromosome-4 with ChromoMap (Anand and Rodriguez Lopez 2022). tRNA annotations were obtained from GtRNAdb (Chan and Lowe 2016). Maternal U1 and U6 sequences were obtained from a curated annotation by (Pagano et al. 2020a). Annotations were visualized in IGV (Thorvaldsdottir et al. 2013). Orthologs for Chr4 genes were obtained from ZFIN (Bradford et al. 2022).

## RESULTS

### Differential gene expression in Nadia gonads

Using RNA-seq, we compared gene expression patterns between testes from chromosomally ZZ adult males and ovaries from chromosomally ZW adult females in 3-month post-fertilization young adults of the Nadia strain, pooling both gonads from each of five replicate individuals of each sex genotype for ten independent samples. Sequencing resulted in an average of 10.45 million reads per library over the ten libraries. Analysis of testis vs. ovary with DESeq2 (Love et al. 2014) identified 17,052 differentially expressed (DE) genes (q-value cutoff of 0.1 and p-value cutoff of 0.05) from the 25,432 annotated protein-coding genes in the GRCz11 zebrafish genome assembly. A total of 6,557 genes showed ovary-biased expression and substantially more (10,495 genes) showed testis-biased expression (Fig. 1A, Supp. Table 1).

**Figure 1.**
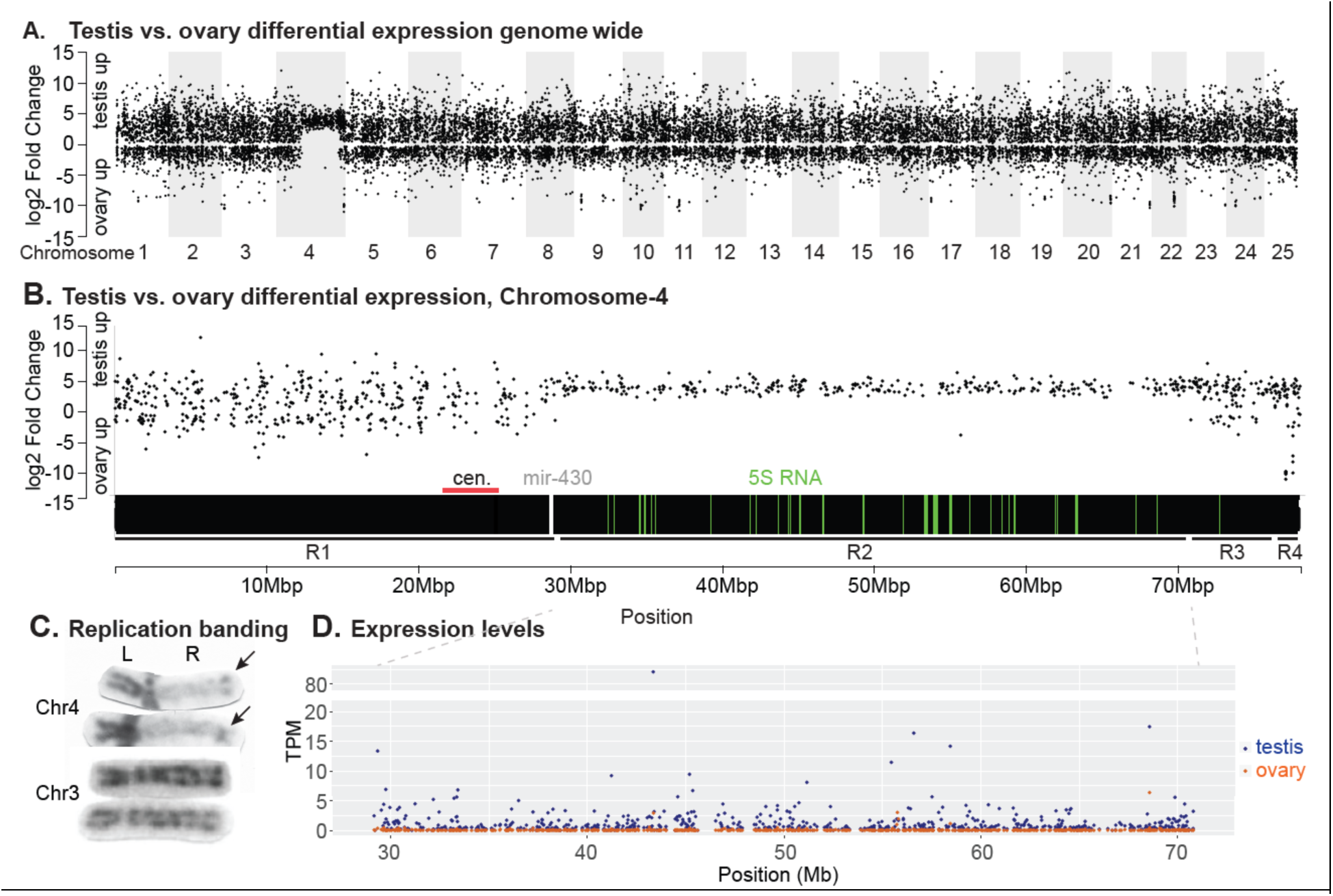
Testis versus ovary differential gene expression. A. Testis versus ovary differential expression plotted against genomic position across the 25 zebrafish chromosomes. The plot shows specific upregulation of testis genes vs. ovary genes on Chr4, the sex chromosome. B. Testis versus ovary differential expression across Chr4.The plot identified four distinct regions: R1, with values matching the bulk of the genome; R2, with testis expression on average much higher than ovary expression; R3, again with values similar to most of the genome; and R4, with some highly ovary-biased genes and coincident with *sar4* containing the major sex determining locus. The horizontal black bar represents Chr4, with the centromere (horizontal red line) located by alignment of the centromeric markers BX537156 (Freeman et al. 2007) and Z20450 (Mohideen et al. 2000) to GRCz11; the positions of maternal 5S ribosomal genes (Locati et al. 2017a) are indicated by green vertical lines, and *mir430* genes are marked by white vertical lines. C. Replication banding of Chr4 and Chr3 (the third and fourth longest chromosomes cytogenetically, respectively (from (Amores and Postlethwait 1999) see also (Phillips et al. 2006)). Karyotypes showed dark staining (early replicating DNA) on the short (left) arm of Chr4 and at the distal tip of the long (right) arm of Chr4, in contrast to pale staining (late replicating DNA) in the position expected for R2 on Chr4R, while only short light bands appeared on all other chromosomes, like Chr3. D. The absolute level of gene expression in transcripts per kilobase of gene length per million total reads (TPM) for all protein coding genes within Region-2 of Chr4, showing that transcripts were detected from most genes in testis (blue) but no transcripts were detected from most genes in ovary (red).

### Functional Enrichment Analysis

To analyze functions of the most strongly differentially expressed genes, we set criteria of greater than 5-fold change and a q-value of less than 0.0001, thereby identifying a subgroup of 6,305 strongly DE genes. Of these strongly DE genes, 1,521 (24.1%) showed ovary-biased expression but substantially more, 4,784 (75.9%), showed testis-biased expression (Fig. 1A, Supp. Table 1). GO term enrichment analysis for Biological Process and Molecular Function for these strongly DE genes identified the top categories for Biological Process classification as “sperm-egg recognition” (GO:0035036); “cell-cell recognition” (GO:0009988); “fertilization” (GO:0009566); “binding of sperm to zona pellucida” (GO:0007339); “oogenesis’ (GO:0048477), “axonemal dynein complex assembly” (GO:0035082), “cilium movement” (GO:0003341); and “germ cell development” (GO:0007281). (Supp. Table 2). For the Molecular Function classification, top categories were “protein serine/threonine kinase activity” (GO:0004674) and ‘protein kinase activity” (GO:0004672). (Supp. Table 3). These categories are expected for germ cell development and cell signaling during gametogenesis.

### Genomic Distribution of DE genes

To determine whether sex bias in differential gene expression was evenly distributed across the genome, we plotted the log2-fold expression change of all 17,052 protein-coding genes that were differentially expressed between adult ZW ovaries and ZZ testes (Fig. 1A). Results showed that for most of the genome, genes with testis-biased expression tended on average to be differentially expressed to a greater fold than genes with ovary-biased expression, confirming earlier studies for zebrafish and other animals, including mammals (Parisi et al. 2003; Ranz et al. 2003; Yang et al. 2006; Malone et al. 2006). On average, testis overexpressed genes were overexpressed 9.2-fold but ovary overexpressed genes were overexpressed only 3.7-fold.

Differentially expressed genes were approximately evenly distributed across the genome, with one exception (Fig. 1A). The sex chromosome, chromosome-4 (Chr4) (Anderson et al. 2012; Wilson et al. 2014) stood out due to a large block of genes substantially upregulated in testis relative to ovary (Fig. 1A).

To examine Chr4 in more detail, we first identified the position of its centromere by aligning the centromeric markers BX537156 and Z20450 (Mohideen et al. 2000; Freeman et al. 2007) to GRCz11 (Fig. 1B). A focus on Chr4 showed that its right arm (Chr4R) in Ensembl (https://www.ensembl.org/Danio_rerio/Location/Chromosome?r=4), which is the cytogenetically long arm, Chr4q (Postlethwait et al. 1994; Daga et al. 1996; Phillips et al. 2006)) had: 1) few genes with ovary-biased expression; 2) an average expression bias for testis-biased genes stronger than the genome-wide average; and 3) less variation in dynamic expression range compared to the left arm (Chr4L, the cytogenetic short arm Chr4p), which was similar to the rest of the genome (Fig. 1A, B). These results show that Chr4 is not only cytogenetically distinct (Pijnacker and Ferwerda 1995; Daga et al. 1996; Gornung et al. 1997; Amores and Postlethwait 1999; Sola and Gornung 2001; Traut and Winking 2001; Phillips et al. 2006), but at least for for Nadia strain gonads, was also transcriptionally distinct.

Differential gene expression for testis vs. ovary divided Chr4 into four distinct regions (Fig. 1B). <u>Region-1</u> extended from the telomere of Chr4L, through the centromere, and ended at position 29.1Mb, just right of the *mir430* gene cluster on Chr4R at position 29.1Mb. Differential gene expression in Region-1 exhibited a pattern like the rest of the genome, with many genes upregulated in the testis and somewhat fewer genes upregulated in the ovary (Fig. 1B). In general, as in most other chromosomes, expression levels in Region-1 tended to be upregulated more in the testis than in the ovary, on average.

<u>Region-2</u> began shortly to the right of the *mir430* gene cluster at 29.1Mb on Chr4R and extended to 70.8M (Fig. 1B). In Region-2, 307 of 308 differentially expressed genes annotated as protein coding were overexpressed in testis compared to ovary (Fig. 1B). These upregulated protein-coding genes overlapped with the positions of maternal 5S ribosomal genes (Locati et al. 2017a) (Fig. 1B). Region-2 of Chr4 appears to be cytogenetically distinct: Chr4R is largely heterochromatic and previously published replication banding (Amores and Postlethwait 1999) showed most of it to be late replicating, in contrast to the right telomeric region and Chr4L, which matched most of the rest of the genome as being early replicating (Fig1C). Because heterochromatin is associated with transcriptional silencing, we wanted to see how Region-2 compared to the rest of the genome. We examined the absolute levels of gene expression across the genome by calculating the TPM for all protein-coding genes in the dataset. We found that for Nadia gonads, the TPM for most genes in Region-2 was zero or nearly so in the ovary (Fig. 1D). The average expression for all 651 annotated protein-coding genes in Region-2 was 0.06 TPM in ovaries regardless of whether genes were differentially expressed in testis vs. ovary; in contrast, the TPM average genome-wide was 39.3, over 600 fold higher. About half of these 651 Region-2 genes (352 genes, 54.1%) lacked detectable expression in the ovary. In contrast, the average expression level for these 651 genes in testes was 1.22 TPM, with only 79 (12.1%) lacking detectable expression. A total of 74 Region-2 genes lacked detectable expression in both gonads. We conclude that genes in Region-2 of Chr4 in Nadia gonads showed greatly reduced transcript numbers compared to the rest of the genome, consistent with its heterochromatic nature, and that this region was transcriptionally silent in ovary.

<u>Region-3</u> stretched from 70.8Mb to 76.9Mb. In Region-3, the relative expression of testis or ovary-biased genes was again distributed as in the bulk of the genome (Fig. 1B).

<u>Region-4 corresponded to the sex-determining region, *sar4*</u> (Wilson et al. 2014). Region-4 began at 76.9Mb at the right of the *ms4a17a* gene cluster and extended to the telomere, and it contains the zebrafish sex-linked loci (Wilson et al. 2014). Region-4 stood out not only because it contains previously identified strongly sex-linked loci, but also because it had some of the most intensely ovary-biased genes across the entire genome (Fig. 1A, B).

Region-4 in GRCz11 contained 58 annotated protein-coding genes, 16 of which were upregulated in ovary relative to testis in the Nadia analysis, and 28 of which were upregulated in testis (Supp. Table 1). Of the 16 ovary-biased genes in Region-4, nine were overexpressed more than 150-fold (log2 fold change = 7.23) relative to testis and four were expressed more than 1000-fold (log2 fold change = 9.97, Supp. Fig. 1). These genes included the mucin gene *CU467646.3* (ENSDARG00000099076), which belongs to a tandemly expanded *mucin-2-like,* cyprinid-specific gene family (gene tree ENSGT00630000090221) that encodes chorion proteins (Hau et al. 2020). Three of the 16 genes highly upregulated in ovary (ENSDARG00000112275, ENSDARG00000113671, ENSDARG00000115978) are members of a cluster of maternal 45S ribosomal genes that were misannotated as protein-coding genes in Ens92 (Ortega-Recalde et al. 2019; Locati et al. 2017b). None of these ovary-biased genes in the sex-determining region are related to any known sex-determination genes in other species or are predicted to play any specific role in gonad development, although the maternal 45S genes, have been proposed to be intrinsic to sex determination in zebrafish (Ortega-Recalde et al. 2019).

Although some highly ovary-specific genes appear to cluster in Region-4 (Fig. 1B), in total, more differentially expressed protein-coding genes in Region-4 showed testis-biased expression than ovary-biased expression. The highest testis-biased Region-4 gene, however, was only 39-fold overexpressed in testis vs. ovary, compared to the hundreds of fold overexpression for some of the ovary-biased genes.

### Gonadal gene expression in a domesticated laboratory strain

In contrast to NA, the laboratory strain AB lacks strongly sex-linked markers and appears to lack a Z chromosome (Wilson et al. 2014), likely because it was produced by several generations of gynogenesis (Walker-Durchanek 1980) and so is likely chromosomally WW, a genotype that usually makes females but also neomales (Wilson et al. 2014; Valdivieso et al. 2022). We wondered if the testis-vs.-ovary-biased gene expression pattern in NA Chr4 was a peculiarity of the chromosomal sex determination system in NA strain fish, or if it also held for the laboratory AB strain. To answer this question, we analyzed RNA-seq data from ovaries and testes of 8-month-old adult AB wild-type controls (SRA accession PRJNA512103 (Yan et al. 2019)). Analysis of transcription levels from ovaries of four AB females and testes of three AB males revealed that AB and NA share the same pattern of gonad-specific gene expression levels in all four regions of Chr4 (Fig. 2). Region-2 had a mean expression level for all protein-coding genes in AB ovaries of 0.17 TPM, with 370 of 651 genes (56.8%) lacking detectable expression; the mean expression level for AB testis was four times higher than AB ovaries (0.68 TPM), with only 20.4% of genes (133 of 651) showing no expression (Supp. Table 4). The overall pattern for AB gonads was like that obtained for NA gonads. This finding shows that the Nadia chromosomal sex determination mechanism is not necessary for the ovary-specific gene silencing observed for Region-2 of Chr4, but instead this silencing must be related to gonad development and/or function.

**Figure 2.**
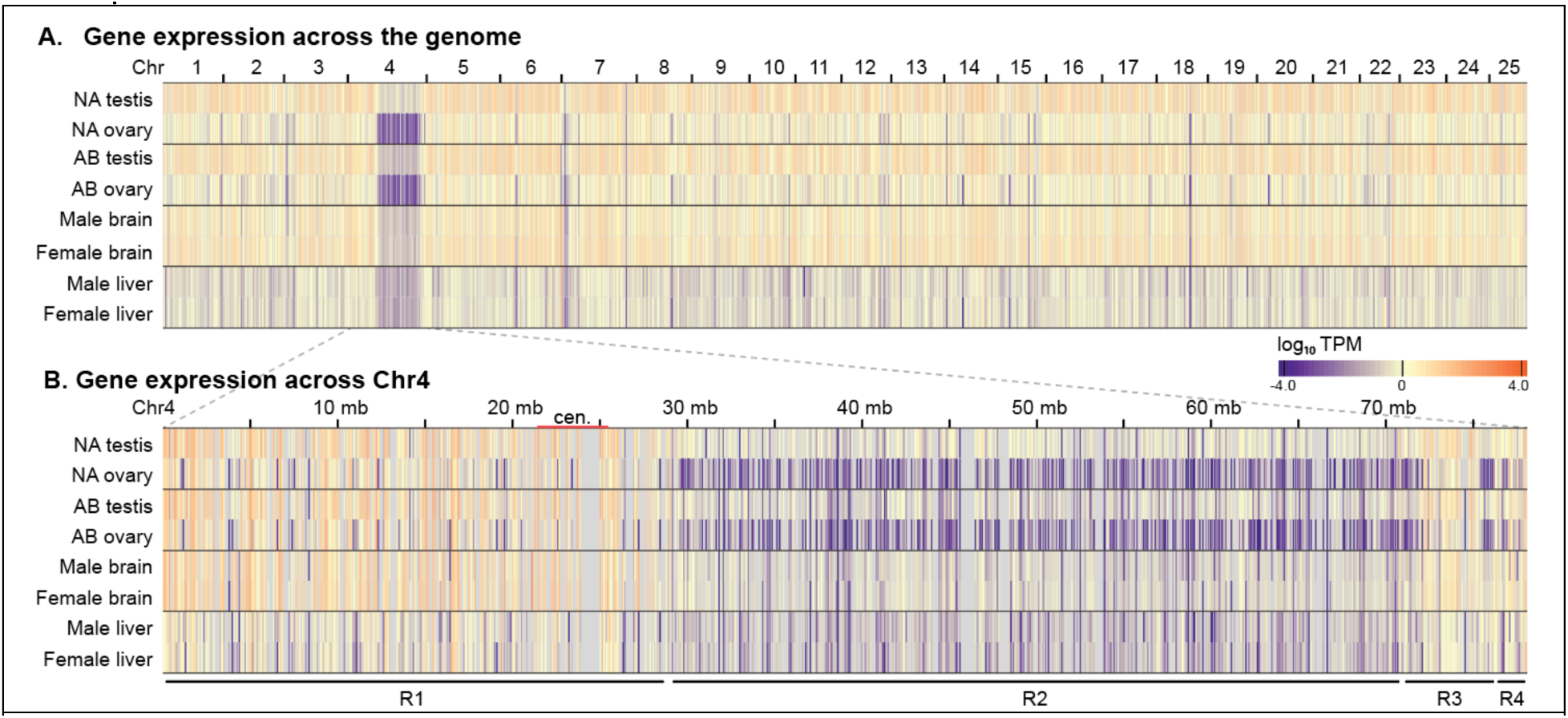
Gene expression in gonads, brain, and liver measured by log_10_ TPM. A. A heatmap showing TPM levels of protein-coding genes across the genome in male and female gonads of strain NA and AB and in male and female somatic organs (brain and liver) for strain AB plotted against position along the genome. Color scale at right. B. A heatmap showing TPM along the length of Chr4 for the same samples as panel A. Plots revealed low transcript levels in Region-2, especially in ovaries of both Nadia and AB, and that, although gene expression was also low in Region-2 in strain AB brain and liver, it was not sexually dimorphic in these somatic organs.

### Expression of Chr4 Region-2 genes in somatic organs

Having shown that ovary-specific silencing of Region-2 depends on testis vs. ovary function and not on sex chromosome genotype, we wondered if the sex-specific expression differences we detected were displayed only by gonads or also occurred in somatic organs. To answer this question, we accessed published RNA-seq data for two additional organs, brain and liver. Data were retrieved from brains of two AB males and three AB females and from livers of two AB males and two AB females, with data from all four samples derived in the same study (PRJNA600413) (Banu et al. 2020). We calculated TPM for all annotated protein-coding genes across the genome and plotted results as for the gonad experiments. Results revealed reduced expression of Chr4 Region-2 genes relative to other portions of the genome for both brain and liver (Fig. 2; Supp. Table 4), paralleling results for gonads (Fig. 1). This finding shows that reduced expression of loci in Region-2 was not specific to gonads but was shared by at least two other somatic organs. A close look at transcript counts revealed low but detectable levels of transcripts of Region-2 genes in brain and liver, unlike ovary, where hardly any transcripts were detected for genes in Region-2. In male livers, the average TPM for genes in Region-2 was 0.27 with 123 of 651 genes (18.9%) lacking detectable expression, and in females, the average TPM for genes in Region-2 was 0.25 with 141 genes (21.7%) lacking detectable expression. In male brains, the average TPM for genes in Region-2 was 0.49 and in females it was 0.57, with 74 (11.4%) and 76 (11.7%) genes lacking detectable expression, respectively (Supp. Table 4). This suppression of gene expression is consistent with the enrichment of transcription-inhibiting H3K9me2 and H3K9me3 histone modifications observed along Chr4R (Yang et al. 2020; Padeken et al. 2022). While repression of transcription for Region-2 genes occurred in both somatic organs we examined, it was not sex-specific in brain or liver, and neither somatic organ showed the apparent silencing of transcription observed for the ovary. We conclude first, that gene expression for Chr4 Region-2 is reduced both for adult gonads and adult somatic organs, and second, that expression is uniquely silent in adult ovaries, and merely suppressed in testes and somatic organs.

### Gene content of Region-2

Chr4 in zebrafish has long been known to be unique in the genome due to its heterochromatic long arm Chr4R (Pijnacker and Ferwerda 1995; Daga et al. 1996; Gornung et al. 1997; Amores and Postlethwait 1999),. The genic content of Chr4R was also shown to be distinct in terms of 5S rRNA genes, *mir430* genes, zinc-finger protein genes, snRNA genes, tRNA genes, Kolobok-1 DNA transposons, and pseudogenes (Anderson et al. 2012). The Chr4R-linked 5S rRNA genes are a maternal-specific isoform (Locati et al. 2017a). The satellite repeats MOSAT-2 and SAT-2 have a complementary distribution, with MOSAT-2 shown to be almost exclusively on Chr4R but SAT-2 being absent from Chr4R although present broadly otherwise throughout the genome (Howe et al. 2013). Chr4R also contains unique gene families that encode mainly either NOD-like receptor (*nlr*) proteins, or zinc finger proteins (*znf*) (Howe et al. 2013; Howe et al. 2016). While unusual features had been ascribed to the entire long arm of Chr4 in earlier versions of the genome assembly, we wanted to understand the relationship of these repetitive genes and satellite DNAs to the four specific regions on Chr4 defined by testis-vs.-ovary gene expression, and so plotted the positions of these features on the GRCz11 assembly (Fig. 3).

**Figure 3.**
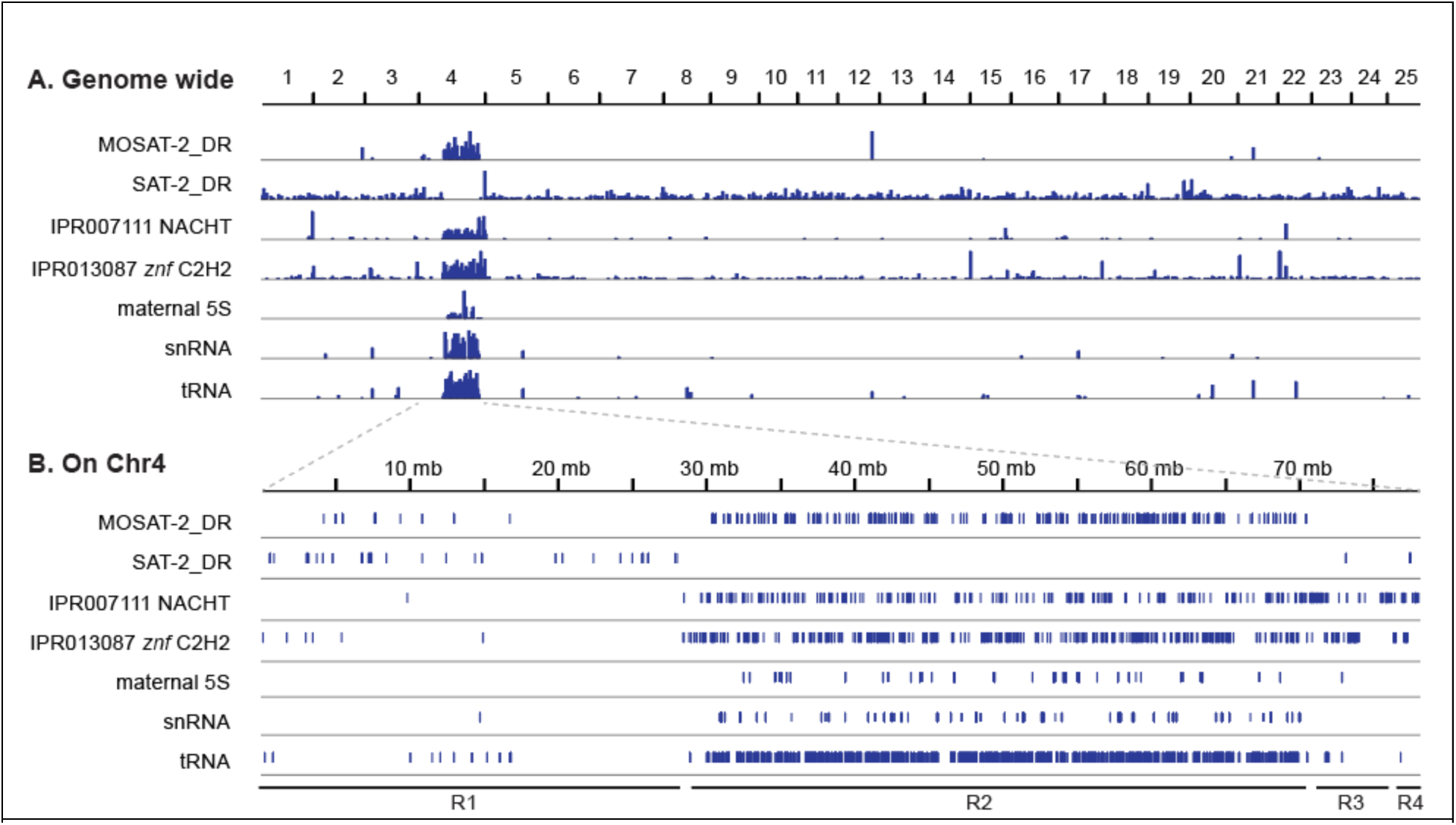
Distinctive features of Chr4. A. Distribution of sequences across the 25 zebrafish chromosomes for satellite sequences MOSAT-2 and SAT2, genes encoding the protein-coding domains IPR007111 NACHT, genes encoding IPR013087 C2H2-type zinc fingers, and certain non-protein-coding genes including maternal 5S rRNA genes (somatic 5S rRNA genes on Chr18 not shown), snRNA genes, and tRNA genes. B. Positions of these sequences along Chr4 are specifically enriched or excluded from Region-2 (Anderson et al. 2012; Howe et al. 2013; Howe et al. 2016). MOSAT-2 is largely restricted to Region-2, while reciprocally, SAT-2 is absent from Region-2. Maternal 5S genes, snRNAs and tRNAs are largely restricted to Region-2, while NACHT-domain containing NLR genes and zinc finger genes are distributed along the length of Chr4R. Although several of these factors had been reported to be specific for Chr4R, we show here that, except for the NACHT and ZNF families, they are restricted to Region-2.

Results showed that the unique features of Chr4 are not evenly distributed along the chromosome. Strikingly, MOSAT-2 distribution was strongly located within Region-2 but was greatly reduced in Region-1 and absent from Regions-3 and −4 and most of the rest of the genome (Fig. 3). Reciprocally, SAT-2 was entirely absent from Region-2, but was again detected in Regions-1, 3 and −4 and across most of the rest of the genome (Fig. 3). Genes encoding NLR proteins (InterPro domain IPR007111) and C2H2 zinc finger proteins (InterPro domain IPR013087) extended beyond Region-2 all the way to the end of the chromosome (Fig. 3). 5S ribosomal genes are found in only two places in the genome (Locati et al. 2017a): Chr4 Region-2 5S rRNA genes (Fig. 3) encode maternal-specific ribosomal components in oocytes that are replaced during development with somatic 5S sequences encoded by 12 somatic 5S rRNA genes on Chr18 (Locati et al. 2017a). Chr4 contains large numbers of snRNA genes (Anderson et al. 2012), nearly all in Region-2 (Fig. 3), that are maternal-specific U1 and U6 spliceosome components, while the two U4 sequences located in Region-1 are somatic (Pagano et al. 2020a). It is not yet known whether the tRNAs on Chr4 (Anderson et al. 2012), accumulated in Region-2, are maternal or somatic, but it is tempting to speculate that they, too, may be components of maternal-specific translation machinery different from somatic tRNA genes. The presence of maternal 5S rRNA and snRNA genes in Region-2 is interesting because these genes are highly expressed in developing oocytes but their position overlaps with protein-coding genes, which conversely, are silenced or nearly so. These observations suggest a reciprocal relationship in Region-2 in ovaries between strong expression of these RNA genes and the silencing of expression for protein-coding genes. The 5S rRNA genes are transcribed by RNA polymerase III, as is U6 (Kunkel et al. 1986; Weinmann et al. 1974), but U1 and protein-coding genes are both transcribed by RNA polymerase II (Weil et al. 1979; Murphy et al. 1982). The expression of maternal U1 sequences from Region-2 suggests that genes in this section of the genome are not globally inaccessible to RNA polymerase II in oocytes and gene silencing in this region must occur through a more nuanced mechanism.

About 80% of the genes on Chr4R lack a human ortholog (Howe et al. 2013). Analysis of orthologs annotated in ZFIN (Bradford et al. 2022) show that this phenomenon is more pronounced in Region-2 than in Regions-3 and −4 (Supp. Table 5). In the euchromatic portion of Chr4R (Region-3 and Region-4 combined), 17 of 221 protein-coding genes (7.7%) had human orthologs, compared to 450 of 603 (74.6%) of genes with human orthologs in Region-1. None of the 651 protein-coding genes in Region-2 had a unique identified human ortholog.

Within Region-2, 177 of 651 genes (27.2%) contained the IPR007111 NACHT_ NTPase domain found in NLR genes (Howe et al. 2016) (Fig. 3). An additional 24 genes lacked the IPR007111 NACHT_ NTPase domain but contained other domains also found in NLR genes, including the IPR003877 SPRY_dom domain and the IPR032675 LRR_dom_sf domain (Howe et al. 2016) (Supp. Table 6), so these 24 genes are also likely members of the NLR family. The functions of this gene family have not been extensively studied in zebrafish, but NLR proteins generally play a role in innate immunity (Laing et al. 2008; Forn-Cuni et al. 2019; Li et al. 2017). Three genes within Region-2 contain the interleukin receptor domain IPR032356 IL17R_fnIII_D1. Together with the NLR genes, the clustering of these genes in Region-2 suggests a possible role in innate immunity.

Nearly half of Region-2 genes (321/651) were annotated as possessing the IPR013087 Znf_C2H2_type zinc finger domain (Fig. 3). Importantly, these Region-2 zinc finger genes are among the first genes to be transcribed during the onset of zygotic genome activation (White et al. 2017).

A number of Region-2 genes have protein domains implicated in chromatin modification. Four genes (ENSDARG00000104681, ENSDARG00000076160, ENSDARG00000115416, ENSDARG00000103283) contain the SET domain IPR001214 and are predicted to have histone methyltransferase activity (Jenuwein et al. 1998). Some of these histone methyltransferases share expression profiles with Chr4R zinc finger genes (White et al. 2017). These genes are members of the gene tree ENSGT00540000072423, which also contains several other genes in Region-2 that are not annotated with the SET domain. Two genes, ENSDARG00000077266 and ENSDARG00000104852, contain the IPR000953 Chromo/chromo shadow domain and are therefore predicted to play a role in chromatin modification (Koonin et al. 1995). Four zebrafish-specific genes in Region-2 (ENSDARG00000096491, ENSDARG00000101046, ENSDARG00000102659, ENSDARG00000101912) contain domain IPR032071, which corresponds to a “domain of unknown function”, DUF4806, which is a subtype of BEN DNA-binding domain (Pan et al. 2023). BEN domains are predicted to recruit chromatin-modifying factors (Abhiman et al. 2008), and it is possible that proteins encoded by these Chr4 Region-2 genes may do so as well. Expression of these chromatin modification factors at the mid-blastula transition might help in the remodeling of chromatin during zygotic gene activation.

Three Region-2 genes (ENSDARG00000090355, ENSDARG00000087265, ENSDARG00000101392) contain the IPR017348 PIM1/2/3 domain of protein serine/threonine kinases, and three additional Region-2 genes (ENSDARG00000094500, ENSDARG00000113519, ENSDARG00000099337) contain the IPR001810 F-box domain, a protein-protein interaction motif.

### Region-2 has a unique repertoire of repetitive elements

The right arm of Chr4 has been reported to be rich in repetitive sequences and young transposable elements compared to the rest of the genome (Anderson et al. 2012; Howe et al. 2013; Chang et al. 2022). We wondered if this was a property of the whole of Chr4R or was specific to Region-2 like MOSAT-2 (Fig. 3B). To find out, we obtained the UCSC genome browser repeat annotation library (Navarro Gonzalez et al. 2021) and compared the repeat content of five different portions of the genome: each of the four Regions of Chr4, and the rest of the genome (Chr1-3 plus Chr5-25). We calculated the percent of each region covered by each repeat element. Hierarchical clustering showed that Region-2 was a clear outlier from the rest of the genome in terms of repetitive element content (Fig. 4A). In contrast, Region-1 was similar in repeat composition to the other 24 chromosomes in the genome, while Region-3 and Region-4 were similar to each other but distinct from the bulk of the genome and quite different from Region-2 (Fig. 4A).

**Figure 4.**
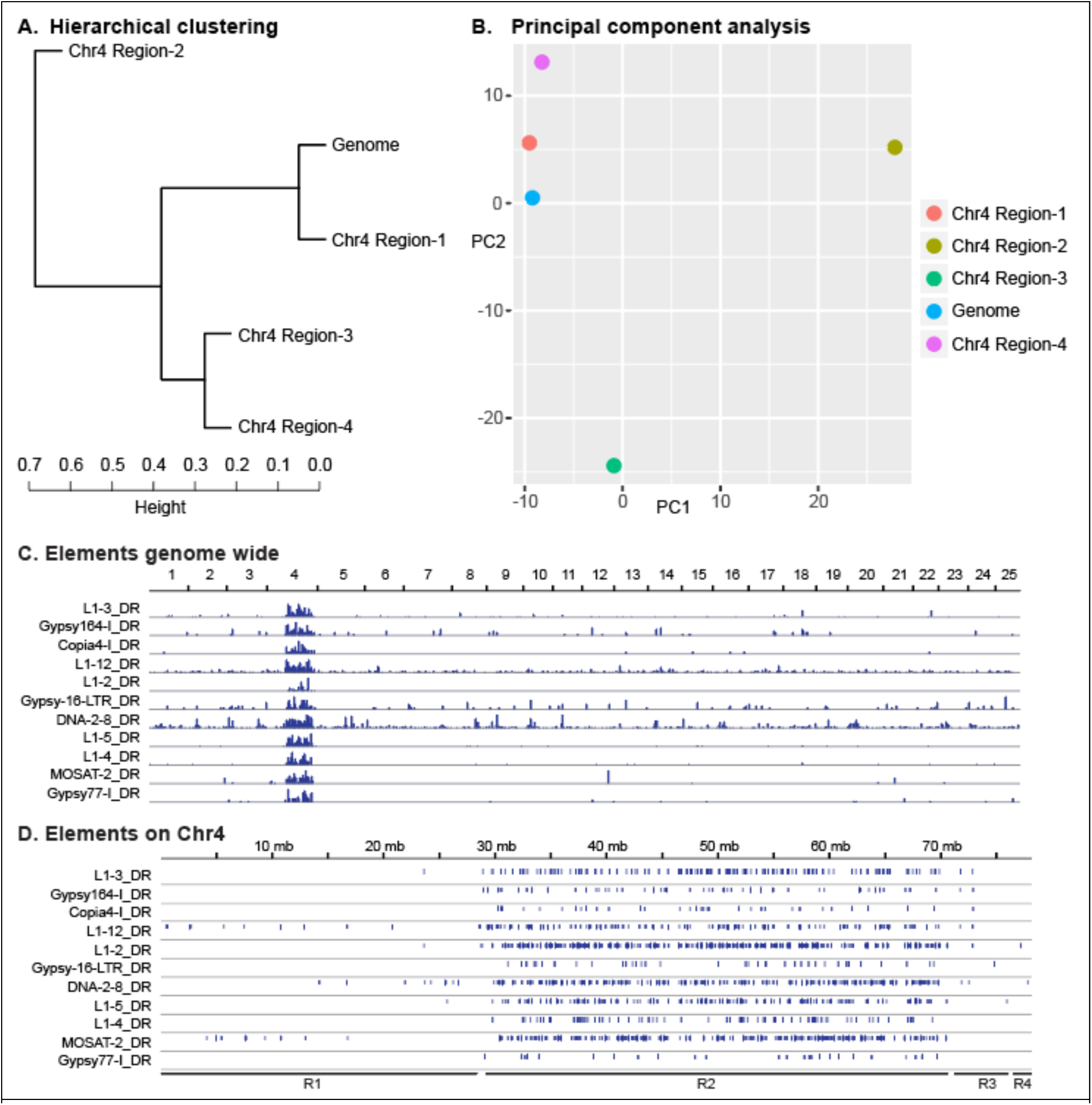
Distribution of repetitive elements. A. Hierarchical clustering analysis of repeat composition of the four regions of Chr4 and the rest of the zebrafish genome. B. PCA analysis of repeat composition shows that Region-2 of Chr4 is distinct from the rest of the genome. C. Genomic distribution of several repeat elements shows them to be clustered on Chr4R. D. Distribution of these repeats along Chr4 shows them to be concentrated in Region-2.

To further explore the distribution of repetitive elements, we conducted a principal component analysis (PCA) of repeat composition for the same five genomic regions. Results showed that PC1 and PC2 explained most of the variance (35.5% and 28.8%, respectively). Region-2 was clearly separated from the rest of the genome along the PC1 axis (Fig. 4B). Examination of the top 25 repeat elements populating PC1 (Supp. Fig. 1) identified several repeat elements highly specific to Region-2 besides MOSAT-2, including DNA-2-8_DR, the LINE elements L1-2_DR, L1-3_DR, L1-4_DR, L1-5_DR, and L1-12 _DR, Copia4-I_DR, Gypsy164-I_DR, Gypsy16-LTR_DR, and Gypsy77-I_DR (Fig. 4C, D). These results show that the uniqueness of Chr4R in terms of its repetitive element content is centered on Region-2.

We wanted to understand if Region-2 of Chr4 exhibited sex-specific transposon expression. We aligned the Nadia gonad RNA-seq dataset to the zebrafish genome and used Telescope (Bendall et al. 2019) to quantify transcript counts for each transposon locus. Results identified 20,509 differentially expressed TE insertions across the genome. Of these, 19,004 loci were upregulated in testis compared to ovary, while only 1,505 loci were upregulated in ovary relative to testis (Supp. Table 7). These results are consistent with published results in medaka, which also found more TE expression in testis (Dechaud et al. 2021). Plotting differentially expressed TEs across the genome showed that Chr4R Region-2 was similar to the rest of the genome in terms of testis-vs.-ovary TE-gene expression despite its uniqueness in terms of TE content (Supp. Fig. 2).

### Chr4 NLR and Zinc Finger genes exhibit different expression patterns

NLR and zinc finger genes occupy Regions-2, −3, and −4 and so we wondered whether the expression of members of these two protein-coding gene families differed according to position along the chromosome. Using the Nadia dataset, we calculated the mean TPM for each NLR family gene (IPR007111 NACHT domain) or zinc finger family gene (IPR013087 C2H2 znf domain). A single Region-2 gene that was annotated with both domains (ENSDARG00000090160) was excluded from analysis. Comparing TPM values of zinc finger genes located in Region-2 vs. Region-3+Region-4 revealed significantly higher expression levels of genes located in Region-3+Region-4 than those in Region-2 for both ovary (p = 2.20E-30) and testis (Wilcoxon rank sums test, p = 4.36E-25) (Fig. 5 A, B). This finding suggests that zinc finger gene family members in Region-3+Region-4 have a regulatory mechanism separate from those in Region-2, and thus, might have a different function. Comparing TPM values of NLR genes in the heterochromatic Region-2 to NLR genes in the euchromatic (Region-3 and Region-4 combined) identified no significant difference between these two regions in either ovary (p = 0.597) or testis (Wilcoxon rank sums test, p = 0.078), although expression of NLR genes in Region-2 and Region-3+Region-4 was somewhat higher in testis than ovary (Fig. 5 A, B). These data further suggest that the NLR family may be regulated by mechanisms different from those regulating zinc finger gene expression, and *nlr* gene silencing along the full-length of Chr4R in ovary suggests that the Nlr proteins may be deleterious to the function of either the ovary or the zygote if they would be provided as maternal transcripts. In the two somatic organs we examined (liver and brain), the zinc finger genes and the NLR genes on Chr4R are significantly more highly expressed in Region-3+Region-4 than in Region-2 (Supp. Fig. 3).

**Figure 5.**
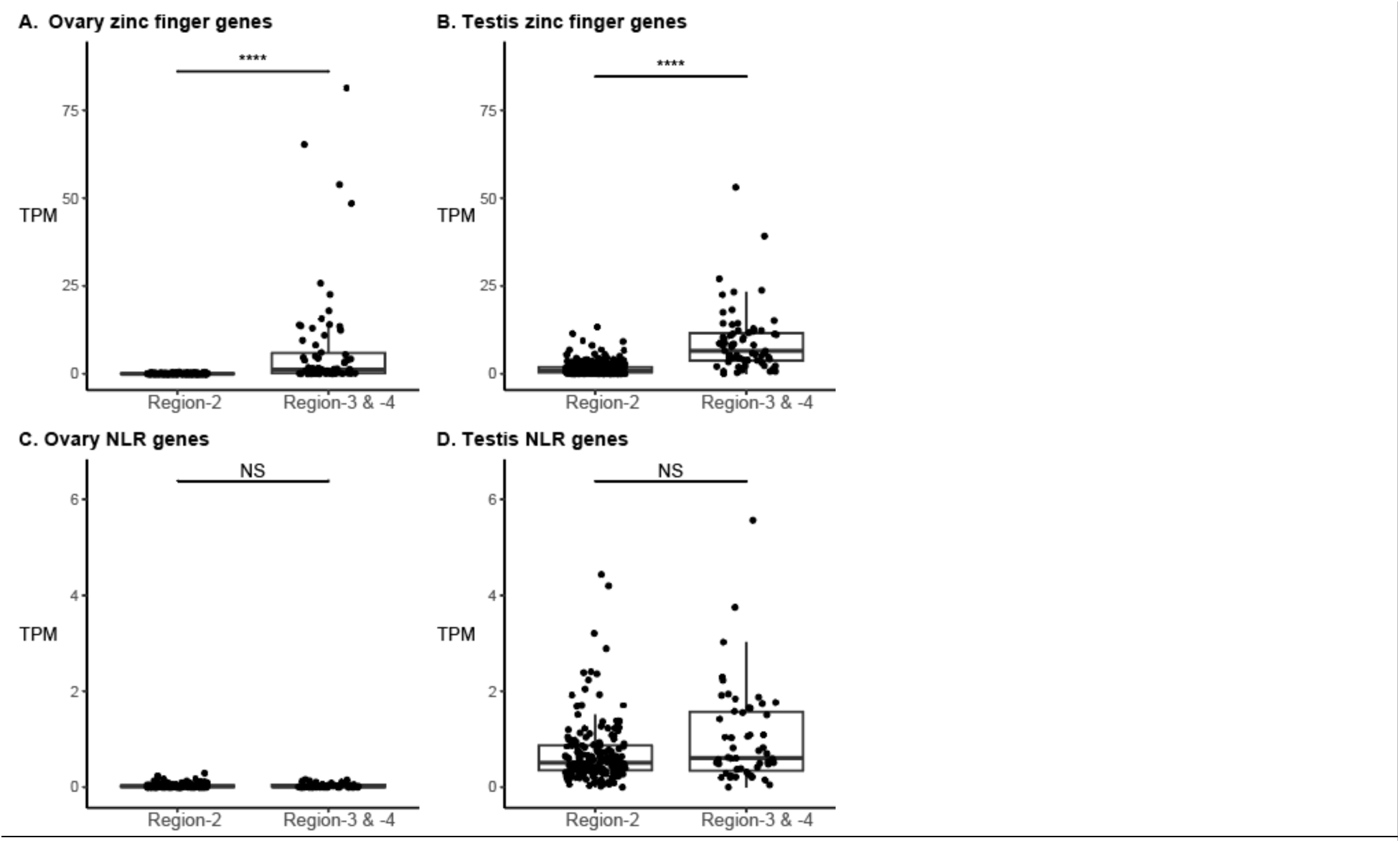
Comparison of expression levels of zinc-finger and NLR genes in Region-2 vs. Region-3+Region-4. A. In the ovary, *znf* genes in Region-2 were not expressed but *znf* genes in Region-3+Region-4 were expressed. B. In the testis, *znf* genes in Region-2 were expressed weakly and *znf* genes in Region-3+Region-4 were expressed more strongly. C. In the ovary, *nlr* genes were nearly silent in both Region-2 and Region-3+Region-4. D. In the testis, *nlr* genes in Region-2 were expressed about as much as *nlr* genes in Region-3+Region-4. Significance levels: NS, not significant (p > 0.05); ****, significant at p < 0.0001.

## DISCUSSION

### A chromosome domain with protein-coding genes specifically silenced in zebrafish ovaries

Well-differentiated sex chromosomes can have properties that are strikingly different from the rest of the genome (Schartl et al. 2016). Sex chromosomes can accumulate genes with sex-specific functions in addition to the sex locus, as in mammals and drosophila (Lahn and Page 1997; Zhang et al. 2020), although not in birds (Xu and Zhou 2020). To explore the properties of sex chromosomes in zebrafish, we conducted RNA-seq experiments in the Nadia strain, which was not manipulated for utility in mutagenesis protocols, in contrast to the two commonly used “wild-type” strains AB and TU (Walker-Durchanek 1980; Streisinger et al. 1981; Mullins et al. 1994). NA and several other unmanipulated zebrafish strains have a ZZ / ZW sex chromosome system with Chr4R as the sex chromosome (Anderson et al. 2006; Wilson et al. 2014). In the NA strain, ZZ fish are always male and ZW fish are usually female but occasionally become neomales, showing that the W is necessary, but not sufficient, for female development (Wilson et al. 2014; Valdivieso et al. 2022).

The RNA-seq experiments reported here defined several regions of Chr4 with respect to the relative expression of protein-coding genes in ZZ testis vs. ZW ovary. The left arm Chr4L and the centromeric portion of the right arm Chr4R up to just right of the *mir430* gene cluster (Region-1) were like the bulk of the genome, with somewhat testis-biased expression of protein-coding genes on average and a greater dynamic range in testis than ovary (Fig. 1). Chr4 Region-1 is euchromatic in cytogenetics (Pijnacker and Ferwerda 1995; Daga et al. 1996; Amores and Postlethwait 1999) (Fig. 1C). Differential gene expression in the middle portion of the right arm (Region-2), however, was unique across the entire genome: the ratio of protein-coding gene expression in testis vs. ovary was greatly biased towards testis with a reduced dynamic range compared to the rest of the genome (Fig. 1A). Region-2 also corresponded to the heterochromatic block on Chr4R (Fig. 1B, C). To the right of Region-2, in Regions-3 and −4, relative expression ratios for testis vs. ovary were again similar to those of autosomes and corresponded in position to a euchromatic region near the right telomere cytogenetically (Fig. 1B, C). The euchromatic portion of Chr4 adjacent to the right telomere including Region-4 is also distinctive because many of the most strongly overexpressed genes in ovary vs. testis were expressed there (Fig. 1B), and it contains the major sex determining locus in natural zebrafish strains (Wilson et al. 2014).

Chr4R contains the largest late-replicating and heterochromatic segment in the zebrafish genome detected in primary cultures of somatic cells (Pijnacker and Ferwerda 1995; Daga et al. 1996; Amores and Postlethwait 1999), suggesting reduced transcriptional activity. Maintenance of heterochromatin relies on the histone demethylase Kdm2a (Frescas et al. 2008), and zebrafish larvae lacking *kdm2aa* activity have upregulated expression of protein-coding genes on Chr4R (Scahill et al. 2017), indicating a role of heterochromatin in inhibiting gene expression in this region. Gene silencing by heterochromatin in sex chromosomes can be a means of dosage compensation, as for Barr bodies in mammals (Barr and Bertram 1949; Ohno et al. 1959; Lyon 1961). The Chr4R heterochromatin block, however, is unlikely to act in dosage compensation for Region-2 genes because Region-2 is present on both the Z and the W in zebrafish and the genes in Region-2 were silenced in ovaries, are expressed in testes, and were downregulated relative to other parts of the genome in somatic organs in both sexes to about the same degree (Fig. 2).

Region-2 of Chr4 contains hundreds of copies of two protein-coding gene families with virtually no transcripts detected in ovaries but low transcript levels found in testes and found in somatic organs (livers and brains) without sex specificity (Figs. 1, 2). We assume that the lack of transcripts from these genes is due to the silencing of transcription, but we cannot formally rule out degradation of these specific transcripts exclusively in zebrafish ovaries. Many of these genes exhibit a burst of transcriptional activity at the maternal-to-zygotic transition (White et al. 2017). Although Region-2 is silenced for protein-coding genes, it actively expresses oocyte-specific genes for transcript splicing (U1 and U6 snRNAs) and maternal transcripts of RNAs necessary for transcript translation into proteins (5S rRNAs) (Locati et al. 2017a; Pagano et al. 2020a), and it also possesses most of the tRNA genes in the genome (Anderson et al. 2012) (Fig. 3A).

To see if the anomalous transcription pattern of Chr4 Region-2 depends on the ZW / ZZ chromosomal sex mechanism, we tested ovaries and testes of AB-strain fish. AB is a domesticated lab line that is likely chromosomally WW due to its origin by gynogenesis (Walker and Streisinger 1983; Walker-Durchanek 1980) (see also https://zfin.org/action/genotype/view/ZDB-GENO-960809-7). Results verified testis-biased gene expression in gonads from domesticated AB zebrafish. The AB result showed that the testis-biased gene expression of Region-2 does not depend on the ZW / ZZ sex-determining mechanism, but rather on biological differences between ovary and testis.

To find whether the male bias in Region-2 transcription is specific to the gonad or if it also appears in somatic organs, we tested male vs. female brains and livers. Results showed that, relative to most of the genome, expression of Region-2 protein-coding genes was generally depressed in somatic organs like it was in the gonads, but that Region-2 expression depression was not sex-specific in somatic organs, in contrast to sex differences in gonads (Fig. 2). We conclude that male vs. female expression bias in Region-2 is specific to gonad development or function. It is possible that Region-2 genes are important for the functioning of testis but not ovary, or that expression of these genes in ovaries would inhibit oocyte differentiation, or that if these transcripts were maternally supplied in the egg, they would hinder zygote development.

### Chr4R Protein-coding genes

Protein-coding genes located on Chr4R are distinct from the rest of the genome. Chr4R in Ensembl Release 92 has 396 zinc-finger genes (see also (White et al. 2017)) and 234 *nlr* genes (see also (Stein et al. 2007; Howe et al. 2016; White et al. 2017)) that are distributed across Regions-2, −3, and −4 (Fig. 3). Transcripts of *znf* genes in Region-2 did not appear in our ovary samples due either to the lack of transcription or, less likely, targeted transcript degradation. In contrast, transcripts of similar *znf* genes in Regions-3 and −4 did appear in ovaries (Fig. 5A).

These results show first, that it is not the whole of Chr4R that is anomalous with regard to transcriptional regulation, but just Region-2, and second, that control mechanisms in the heterochromatic (Region-2) and euchromatic (Region-3+Region-4) portions of Chr4R are likely different. In contrast to ovaries, transcripts from *znf* genes in Region-2 did appear in testes, but even in testes, more transcripts were found from *znf* genes located in Region-3+Region-4 than in Region-2 (Fig. 5B). This result showed that transcripts for *znf* genes located in the heterochromatic portion of Chr4R were less abundant than transcripts for *znf* genes found in the euchromatic portion of Chr4R in both ovary and testis.

<u>The *znf* genes in</u> Region-2 are interesting not only because of their ovary-specific silencing, but also because of their pattern of expression in zebrafish embryos. These Chr4 *znf* genes initiate expression in concert at the beginning of zygotic transcription, increasing from the 1,000-cell stage (3.0hpf) to dome stage (4.3hpf) and then decreasing by the 75% epiboly stage (8.0hpf) (Kimmel et al. 1995; White et al. 2017). Thus, the Region-2 *znf* genes are 1) specifically silenced in ovaries; 2) expressed at low levels in testes, brains, and livers; 3) transcribed early in the maternal-to-zygotic transition (MZT), and then 4) rapidly degraded. Furthermore zebrafish oocytes lacking function of *kdm2aa,* which is necessary for normal heterochromatin function, overexpress Chr4R *znf* genes, and when mated to wild-type sperm, produce defective eggs and many inviable embryos (Scahill et al. 2017). These data suggest the hypothesis that the silencing of Chr4R *znf* genes in oocytes occurs either because these Znf proteins are harmful to oocyte development (most *kdm2aa* mutants develop as males, a phenotype consistent with oocyte death in the juvenile ovary (Rodriguez-Mari et al. 2010)) or detrimental to embryos if these proteins are present before the normal schedule of zygotic gene expression (Scahill et al. 2017). The difference in expression of the Region-2 *znf* genes and the more telomeric Region-3 and Region-4 *znf* genes suggests that the telomeric zinc finger proteins may have a different regulatory mechanism and function.

<u>The *nlr* genes</u> in Region-2, like the *znf* genes, were silenced in ovaries, but unlike *znf* genes, *nlr* genes in Regions-3 and −4 were also silent in ovaries, and even for testes, the difference in expression levels of *nlr* genes in Region-2 compared to Regions-3 and −4 was not significant (Fig. 5C, D). This result suggests that NLR proteins might be deleterious in ovary. In brain and liver, expression of *nlr* genes in Region-2 was significantly less than for *nlr* genes in Regions-3 and −4, consistent with the heterochromatic inhibition of gene expression in Region-2 in somatic cells (Pijnacker and Ferwerda 1995; Daga et al. 1996; Gornung et al. 1997; Amores and Postlethwait 1999; Sola and Gornung 2001; Traut and Winking 2001; Phillips et al. 2006).

Expression of at least 148 of the *nlr* genes on Chr4R was detected during embryogenesis, but they did not exhibit the same expression pattern as the *znf* genes (White et al. 2017). Thus, although the *nlr* genes of Chr4R are downregulated in ovary vs. testis, in general, they don’t show the burst of transcription around the maternal-to-zygotic transition that their *znf* neighbors show. Thus, despite their interspersed location along Chr4R (Fig. 3B), expression of these *znf* and *nlr* genes must depend on different regulatory mechanisms.

<u>Mucin genes</u> in Region-4 included some of the most strongly overexpressed genes in adult zebrafish ovaries vs. testes in the entire genome (Fig. 1B). The most strongly overexpressed ovary-vs.-testis genes included *CU467646.3*, a member of a tandemly expanded *mucin-2-like* cyprinid-specific gene family (see gene tree ENSGT00630000090221) that encodes oocyte-expressed chorion proteins (Hau et al. 2020). None of the ovary-biased protein-coding genes in Region-4 are related to any known genes in the sex-determination pathway in other species. We conclude that the egg chorion genes located in Region-4 likely represent another case of sex-specific genes captured on a sex chromosome.

### Region-2 and *mir430*

A cluster of more than 50 *mir430* genes lies adjacent to the left border of Region-2 (Nudelman et al. 2018; Desvignes et al. 2022). MicroRNAs can modulate gene expression by binding to messenger RNAs to regulate their translation or degradation (Fabian et al. 2010; Huntzinger and Izaurralde 2011). Mature products of *mir430* genes regulate the clearance of maternal mRNAs from zebrafish embryos by causing deadenylation and hence decay but they are not supplied to the egg during oogenesis (Giraldez et al. 2005; Takacs and Giraldez 2016; Giraldez et al. 2006; Lund et al. 2009). Instead, *mir430* gene expression in zebrafish begins after the 64-cell stage (2hpf) concomitant with the recruitment of the cohesin subunit Rad21 (Meier et al. 2018); *mir430* expression peaks at 4hpf, and then decreases after 24hpf (Heyn et al. 2014; Chen et al. 2005). The bulk of transcription of zygotic genes begins mostly after the 1,000-cell stage at 3hpf (Kane and Kimmel 1993; Lee et al. 2014), and zygotic transcripts replace maternal transcripts. Within the nuclei of early zebrafish embryos, *mir430* forms a transcription compartment with their neighboring *znf* genes (Hadzhiev et al. 2019). The regulatory relationships of *mir430* and *znf* loci in Region-2 in ovaries and their activation around the MZT, however, are not yet fully known.

### Chr4R Region-2: a maternal source for the zygote’s translation machinery

<u>Maternal ncRNA genes</u> are clustered on Chr4R. Vertebrate eggs generally contain large numbers of ribosomes packed into oocytes that aid in translating maternal transcripts before zygotic genes become transcriptionally active (Brown and Gurdon 1964). Zebrafish have distinct maternal and somatic RNA genes (Locati et al. 2017a; Locati et al. 2018; Locati et al. 2017b; Pagano et al. 2020a; Pagano et al. 2020b; Ortega-Recalde et al. 2019). On zebrafish Chr4R, Region-2 has nearly all of the 2,330 5S rRNA genes on Chr4R (Fig. 3B), which are expressed in oocytes but not in somatic cells of adult fish (Locati et al. 2017a), while a locus on Chr18 contains twelve 5S rRNA genes that, reciprocally, are expressed in somatic cells of embryonic and adult fish but not in oocytes (Gornung et al. 2000; Locati et al. 2017a; Tao et al. 2020).

While each of the many 5S rRNA genes in eukaryotic genomes is transcribed into a single 5S rRNA molecule, each of the several 45S rRNA genes is transcribed as a single transcript that is processed to form the 18S, 5.8S, and 28S ribosome components (Long and Dawid 1980). rRNA gene expression was not assayed in our analysis due to selection for polyadenylated transcripts and alignment to annotated protein-coding genes, but our data showed a single gene in Region-2 that was upregulated in ovary vs. testis (ENSDARG00000115819, Fig. 1), and sequence analysis showed that this was an 18S ribosomal gene that was misannotated as a protein-coding gene in Ensembl. This result shows that Region-2 has at least one 18S gene that is expressed maternally. In addition, three 45S genes misannotated as protein-coding genes (ENSDARG00000112275, ENSDARG00000113671, ENSDARG00000115978) in Chr4 Region-4 were overexpressed in ovaries vs. testes in our dataset. These 45S genes are expressed exclusively in oocytes and encode rRNAs for maternally supplied ribosomes (Locati et al. 2017b; Ortega-Recalde et al. 2019). These maternal ribosomes also have a specific pattern of 2′-O-methylation different from somatic ribosomes (Ramachandran et al. 2020). Thus, Region-2 contains at least one maternal 18S gene and Region-4 contains at least three other maternal 45S genes, verifying the roles of Region-2 and other parts of Chr4R in supplying maternal ribosome components, while other genes provide somatic / zygotic ribosomal RNAs (Locati et al. 2017b; Locati et al. 2018; Locati et al. 2017a).

<u>Maternal spliceosome components</u> also occupy Region-2. Genes encoding U1 and U6 spliceosome components that are specifically transcribed in oocytes are in Region-2, making these snRNAs available for splicing either maternal messages in the oocyte or the earliest transcripts in the embryo (Anderson et al. 2012; Pagano et al. 2020a; Pagano et al. 2020b). The location of maternal 5S and spliceosome component genes in Region-2 on the zebrafish sex chromosome represent another example of sex-specific functions encoded on the sex chromosome.

<u>Transfer RNA genes</u> are also mostly concentrated in Region-2 of Chr4 (Anderson et al. 2012) (Fig. 3), although tRNAs have not yet been studied with respect to maternal vs. zygotic / somatic cell expression. tRNA genes are also found elsewhere in the genome, so a hypothesis is that the Region-2 tRNA genes are also maternal-specific and thus contribute to specialized maternal vs. zygotic translation machinery along with Region-2 ribosomal and spliceosomal genes.

### Ch4R transposable elements

Chr4R carries a distinctive array of repetitive elements (Anderson et al. 2012; Howe et al. 2013; Chang et al. 2022), but we found that the overall pattern of differential-expression of transposable elements in testis vs. ovary was not different in Region-2 from the rest of the genome. Thus, for the gonads at least, the heterochromatic nature of Region-2 does not block the expression of several types of genes, including transposable element genes, maternal 5S rRNA genes, and maternal U1 and U6 snRNA genes.

### A Maternal-to-Zygotic-Transition Gene Block

In zebrafish, chromosome-4 plays an important role in gonad development and early embryogenesis (Fig. 6). Region-4 near the right telomere of Chr4 contains three groups of genetic elements important for egg development: 1) a factor that is necessary but not sufficient for the bipotential gonad to develop into an ovary in natural zebrafish strains (Wilson et al. 2014); 2) mucin genes encoding egg chorion proteins (Fig. 1); and 3) several 45S ribosomal RNA genes that are expressed specifically in oocytes but not in somatic cells, which express a different set of 45S rRNA genes (Ortega-Recalde et al. 2019; Locati et al. 2017b).

**Figure 6.**
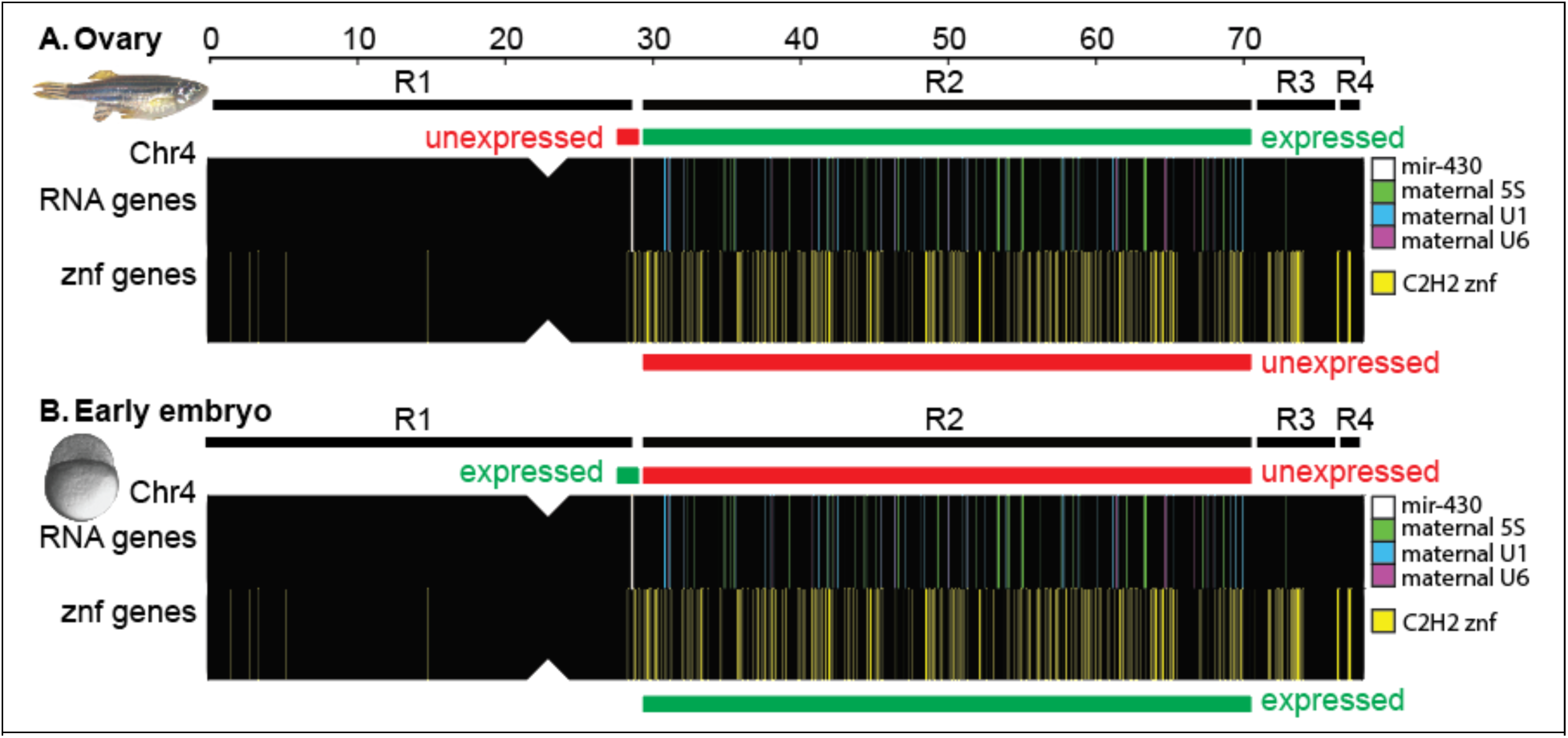
A maternal-to-zygotic transition gene block. A. In the ovary, oocytes express maternal 5S rRNA (green) and spliceosome genes (U1, U6; blue and purple) in Region-2 but not *znf* genes (yellow) in Region-2 or the adjacent *mir430* genes (white) (Locati et al. 2017a; Locati et al. 2018; Locati et al. 2017b; Pagano et al. 2020a; Pagano et al. 2020b). The 5S ribosomal and the spliceosomal maternal transcripts help make the machinery to translate proteins essential for embryogenesis from maternal mRNAs before expression of the zygotic genome. Maternal 45S genes are in Region-4 (Ortega-Recalde et al. 2019). B. Reciprocally, early embryos at the beginning of the maternal-to-zygotic transition do not express maternal 5S rRNA or maternal spliceosome (U1, U6) genes in Region-2, but do express Region-2 *znf* genes and the nearby *mir430* genes. Znf proteins may help to stabilize the mobility of transposable elements (White et al. 2017; Hadzhiev et al. 2019; Wells et al. 2023) and the *mir430* mature products promote clearance of maternal messenger RNAs (Giraldez et al. 2005; Takacs and Giraldez 2016; Giraldez et al. 2006; Lund et al. 2009). Abbreviations: R1, R2, R3, and R4 represent Regions-1 to −4.

Chr4 Region-2 has genes involved in various ways in the production of the maternally inherited gene products that drive the progression of embryonic cell cleavage and the embryo’s transition from maternally derived components to zygotically derived machinery. Maternally expressed 5S rRNA genes and spliceosome factors (U1, U6) lie in Region-2 (Fig. 6A), while somatic alternatives for these RNAs are located elsewhere in the genome (Anderson et al. 2012; Locati et al. 2017a; Pagano et al. 2020a; Pagano et al. 2020b). Ovaries also express at least one 18S rRNA gene, which is usually a part of a 45S rRNA gene (Fig. 1), although most maternal 45S rRNA genes are in Region-4 (Ortega-Recalde et al. 2019). In contrast to ovarian expression of these translation machinery genes (Locati et al. 2017b; Pagano et al. 2020a) (Locati et al. 2017b; Pagano et al. 2020a; Pagano et al. 2020b; Ortega-Recalde et al. 2019), we found expression silencing of protein-coding genes located in Region-2 in ovaries (Fig. 1, 5, 6A).

In embryos, expression patterns of Region-2 genes are reversed from those in ovaries (Fig. 6B). In late cleavage at about the 1000 cell stage (3.0 hours post fertilization, hpf), the rate of cleavage slows, the cell cycle lengthens, and cell divisions become asynchronous in the mid-blastula transition (Kane and Kimmel 1993). About a half hour earlier, however, several *znf* genes in Region-2 begin to be expressed and many burst into transcription from dome stage to 75% epiboly (4.3-8 hpf) (White et al. 2017) (Fig. 6B). The role of at least some proteins encoded by these activated Region-2 *znf* genes appears to be to repress the expression of retroelements during zygotic genome activation (ZGA) (White et al. 2017; Hadzhiev et al. 2019; Wells et al. 2023), and without that suppression, early development goes awry. Autosomal copies of translation machinery genes, including snRNA genes, ribosomal RNA genes, and we predict tRNA genes, become active (Locati et al. 2017a; Locati et al. 2018; Locati et al. 2017b; Pagano et al. 2020a; Pagano et al. 2020b), replace the maternal translation machinery, and translate zygotic mRNAs as the embryo transitions to the use of its own genome. These changes are associated with the redistribution of the cohesion nuclear architecture subunit Rad21 specifically from Region-2 to euchromatic regions of the genome (Meier et al. 2018). At this stage, *mir430* genes near Region-2 become active (Fig. 6B) and *mir430* gene products mediate the decay of maternal mRNA transcripts (Giraldez et al. 2005; Giraldez et al. 2006; Bazzini et al. 2012).

In conclusion, the dramatic lack of ovarian transcripts for Chr4 Region-2 protein coding genes discovered here coupled with the oocyte’s expression of Region-2 maternal translation machinery genes and the subsequent reversal of these patterns associated with zygotic gene activation shows that Chr4 Region-2 is a Maternal-to-Zygotic-Transition Gene Block.

## DATA AVAILABILITY

Sequences are deposited in the Sequence Read Archive under accession PRJNA504448. Supplemental data have been uploaded to the GSA Figshare portal. Supplemental Table 1 lists significantly differentially expressed genes comparing Nadia testis vs. Nadia ovary. Supplemental Table 2 shows Biological Processes associated with highly differentially expressed genes between Nadia testis and ovary (Panther GO-Slim). Supplemental Table 3 shows Molecular Functions associated with highly differentially expressed genes comparing Nadia testis vs. Nadia ovary (Panther GO-Slim). Supplemental Table 4 lists Transcripts per kilobase-million (TPM) values for all annotated protein coding genes in all libraries analyzed (Ens92). Supplemental Table 5 presents Human orthologs for all annotated protein-coding genes on zebrafish Chr4. Supplemental Table 6 displays annotated protein domains for Chr4 Region-2 genes (Ens92). Supplemental Table 7 lists transposable element genes differentially expressed between Nadia testis vs. ovary. Supplemental Figure 1 shows the genomic distribution of the 25 repeat elements contributing most strongly to PC1 in the principal components analysis displayed in Fig. 4, main text. Supplemental Figure 2 compares the expression of transposable element genes in testis vs. ovary. Supplemental Figure 3 compares expression levels for zinc finger genes and NLR genes in the heterochromatic portion (Region-2) and euchromatic portion (Region-3+Region-4) of Chr4R in AB-strain zebrafish.

<u>Supplemental materials</u> available at G3 online.

## ACKNOWLEDGEMENTS

This work benefited from access to the University of Oregon high performance computing cluster, Talapas. We thank our colleagues Angel Amores, Thomas Desvignes, Clay Small, and Yi-lin Yan for helpful conversations.

## SUPPLEMENTAL FIGURES

**Supplemental Figure 1.**
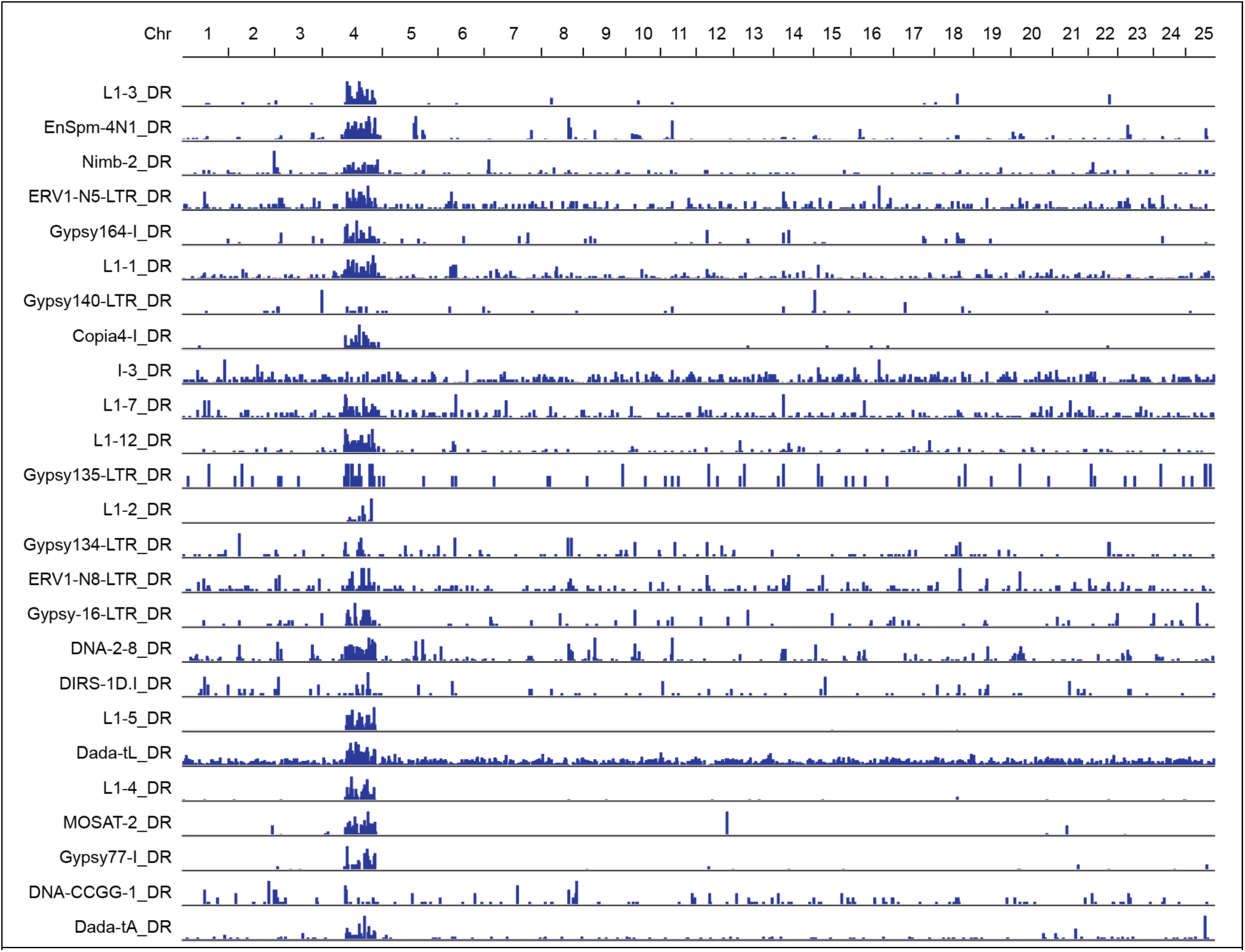
Genomic distribution of the 25 repeat elements contributing most strongly to PC1 in the principal components analysis displayed in Fig. 4, main text.

**Supplemental Figure 2.**
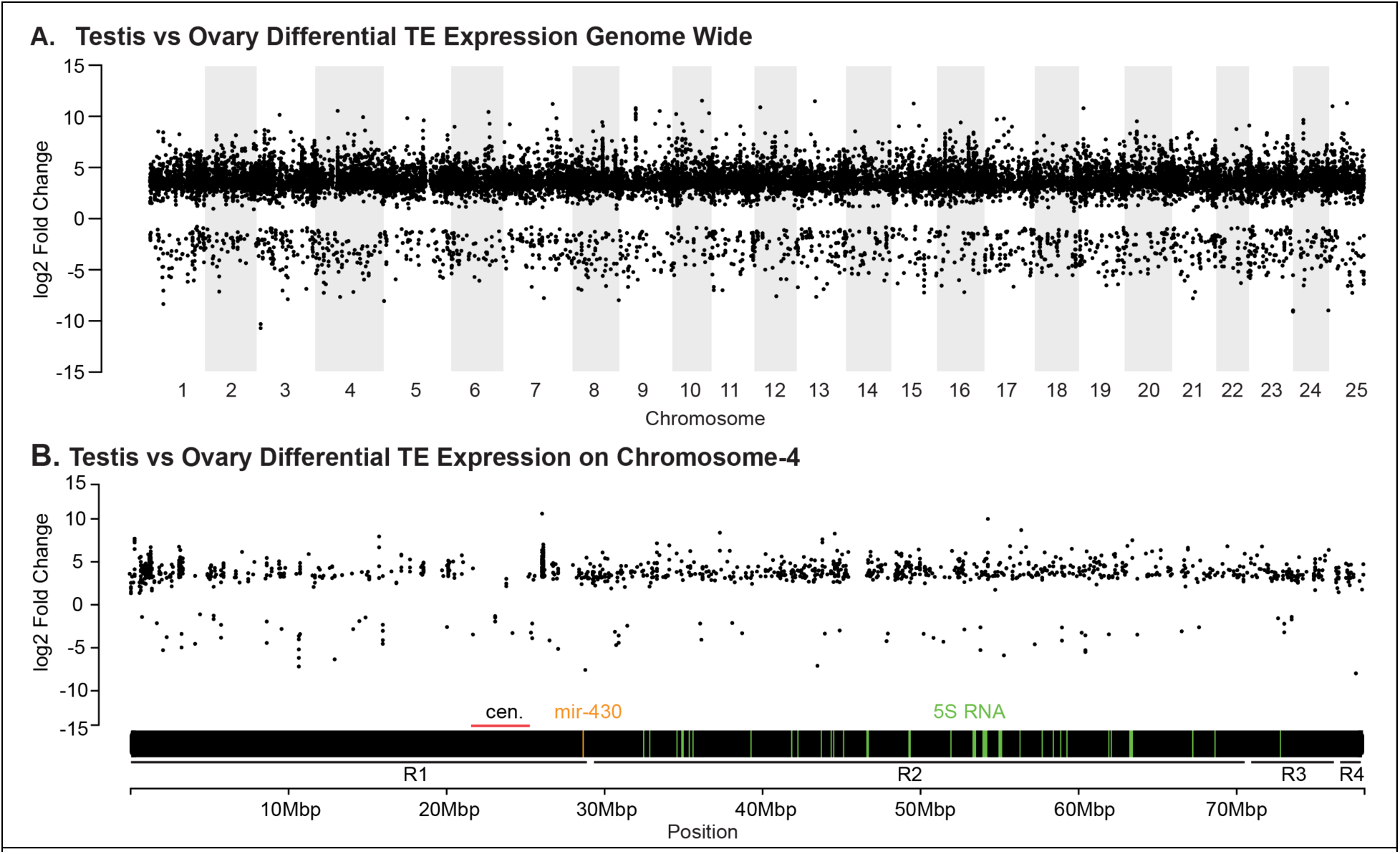
Comparison of the expression of transposable element genes in testis vs. ovary. **A.** Testis versus ovary differential expression of transposable element genes plotted against position genome wide revealed that transposable elements are, in general, more strongly expressed in testis than in ovary and that Chr4R matches the rest of the genome. **B.** Testis versus ovary differential expression of transposable element genes across Chr4 did not show expression silencing in Region-2, in contrast to the result for protein-coding genes shown in Figure 1 of the main text.

**Supplemental Figure 3.**
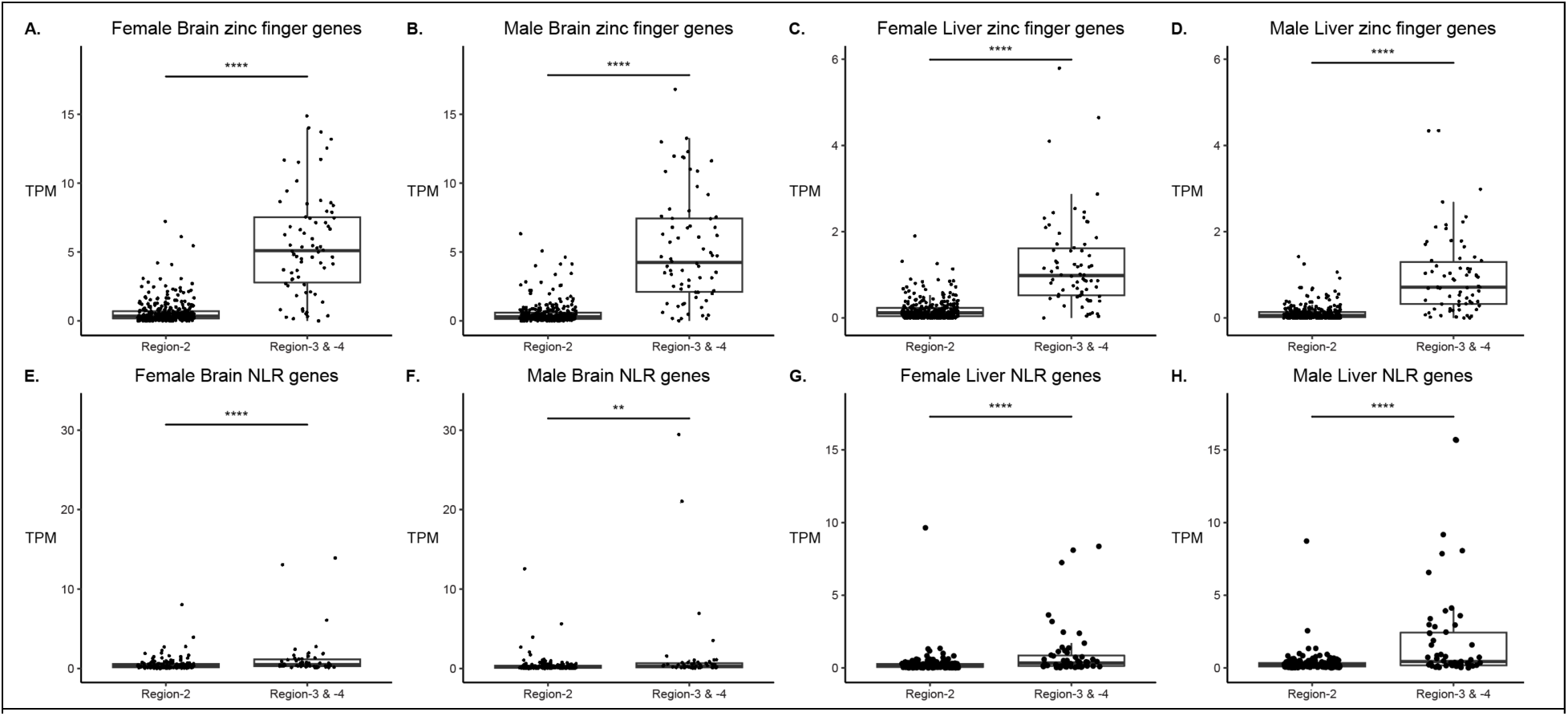
Comparison of expression levels for zinc finger genes and NLR genes in the heterochromatic portion (Region-2) and euchromatic portion (Region-3+Region-4) of Chr4R in AB-strain zebrafish. Results showed significantly higher expression of zinc finger genes in Region-3+Region-4 in (**A**) female and (**B**) male brain and in (**C**) female and (**D**) male livers. This expression pattern was also observed in NLR genes, with Region-2 NLR genes showing lower expression levels than Region-3+Region-4 NLR genes in both brain (**E** and **F**) and liver (**G** and **H**). These results revealed that, like gonads (Fig. 5, main text) these somatic organs also showed depressed expression in Region-2 protein-coding genes.

## SUPPLEMENTAL TABLES

Supplemental Table 1. Significantly differentially expressed genes comparing Nadia testis vs. Nadia ovary.

Supplemental Table 2. Biological processes associated with highly differentially expressed genes between Nadia testis and ovary (Panther GO-Slim). Analysis Type: PANTHER Overrepresentation Test (Released 20230705). Annotation Version and Release Date: PANTHER version 17.0 Released 2022-02-22. Analyzed List: *Danio rerio*. Reference List: panther control list.txt *Danio rerio*. Test Type: FISHER. Correction: FDR.

Supplemental Table 3. Molecular functions associated with highly differentially expressed genes comparing Nadia testis vs. Nadia ovary (Panther GO-Slim). Analysis Type: PANTHER Overrepresentation Test (Released 20230705). Annotation Version and Release Date: PANTHER version 17.0 Released 2022-02-22. Analyzed List: Danio rerio. Reference List: panther control list.txt (Danio rerio). Test Type: FISHER. Correction: FDR.

Supplemental Table 4. Transcripts per kilobase-million (TPM) values for all annotated protein coding genes in all libraries analyzed (Ens92).

Supplemental Table 5. Human orthologs for all annotated protein-coding genes on zebrafish Chr4.

Supplemental Table 6. Annotated protein domains for Chr4 Region-2 genes (Ens92).

Supplemental Table 7. Transposable element genes differentially expressed between Nadia testis vs. ovary.

